# Development of a 3D atlas of the embryonic pancreas for topological and quantitative analysis of heterologous cell interactions

**DOI:** 10.1101/2021.04.28.441857

**Authors:** Laura Glorieux, Aleksandra Sapala, David Willnow, Manon Moulis, Shlomit Edri, Jean-Francois Darrigrand, Anat Schonblum, Lina Sakhneny, Laura Schaumann, Harold F. Gómez, Christine Lang, Lisa Conrad, Fabien Guillemot, Shulamit Levenberg, Limor Landsman, Dagmar Iber, Christophe Pierreux, Francesca M. Spagnoli

## Abstract

Generating comprehensive image maps, while preserving spatial 3D context, is essential to quantitatively assess and locate specific cellular features and cell-cell interactions during organ development. Despite the recent advances in 3D imaging approaches, our current knowledge of the spatial organization of distinct cell types in the embryonic pancreatic tissue is still largely based on 2D histological sections. Here, we present a light-sheet fluorescence microscopy approach to image the pancreas in 3D and map tissue interactions at key development time points in the mouse embryo. We used transgenic mouse models and antibodies to visualize the three main cellular components within the developing pancreas, including epithelial, mesenchymal and endothelial cell populations. We demonstrated the utility of the approach by providing volumetric data, 3D distribution of distinct progenitor populations and quantification of relative cellular abundance within the tissue. Lastly, our image data were combined in an open source online repository (referred to as Pancreas Embryonic Cell Atlas). This image dataset will serve the scientific community by enabling further investigation on pancreas organogenesis but also for devising strategies for the *in vitro* generation of transplantable pancreatic tissue for regenerative therapies.

## Introduction

Organogenesis is a fine-tuned process, involving the expansion and differentiation of organ-specific progenitor cell populations in the context of profound morphological changes in the tissue architecture (Zorn and Wells, 2009). The formation of an organ relies on the precise activation of cell-intrinsic differentiation programs but also on the interaction of progenitor cell populations with surrounding tissues. These cell-cell interactions provide essential cues to guide cell differentiation and tissue morphogenesis in a spatially- and temporally-defined fashion. Full comprehension of these events will help to elucidate the mechanisms underpinning tissue formation but also guide bioengineering strategies to build three-dimensional (3D) tissue models (Bulanova et al., 2017; Iber et al., 2016; Kaufman-Francis et al., 2012).

The pancreas represents a paradigmatic example of how intrinsic cell–cell interactions and extrinsic signals released from surrounding non-pancreatic tissues, such as mesenchyme and blood vessels, coordinate cell differentiation and morphogenesis to form a fully functional adult organ (Cozzitorto and Spagnoli, 2019; Larsen and Grapin-Botton, 2017; Pan and Wright, 2011; Shih et al., 2013; Wessells and Cohen, 1967). The adult pancreas is an amphicrine gland whose exocrine compartment produces and releases the digestive enzymes, while the endocrine compartment houses the insulin-secreting β-cells and is critical for blood glucose homeostasis (Sneddon et al., 2018). Remarkably, all pancreatic cell types, including the acinar, ductal and endocrine cells, derive from a common pool of endoderm progenitors, which is specified in the mouse around embryonic day (E) 8.5 (Gittes, 2009; Zorn and Wells, 2009). Between E12.5 and E14.5, the pancreatic epithelium undergoes important remodeling events, which result in the formation of a tubular network called plexus, with a proximo-distal ‘tip and trunk’ domain organization (Bankaitis et al., 2015; Larsen et al., 2017; Pan and Wright, 2011; Pierreux et al., 2010; Sznurkowska et al., 2018; Villasenor et al., 2010; Zhou et al., 2007). Regions at the periphery of the pancreatic epithelium consist of elongating branch tips, while the center contains the luminal plexus and corresponds to trunk domains. Importantly, such proximo-distal architecture coincides with cell fate restriction of pancreatic progenitors and subsequent differentiation, whereby cells at the “tip” adopt an acinar differentiation program, while cells within the “trunk” remain in a bipotent state and contribute to the ductal and endocrine cell lineages during further development (Fig. 1A’). Neighboring mesenchymal cells support this developmental process and a vascular network forms throughout the growing pancreas (Angelo and Tremblay, 2018; Azizoglu and Cleaver, 2016; Cozzitorto and Spagnoli, 2019; Golosow and Grobstein, 1962; Sakhneny et al., 2019; Seymour and Serup, 2019). Depletion of the mesenchyme impairs pancreatic epithelial growth, morphogenesis as well as differentiation of acinar and β-cells, suggesting a fundamental regulatory role for these cells throughout embryonic development (Attali et al., 2007; Cozzitorto et al., 2020; Esni et al., 2001; Harari et al., 2019; Landsman et al., 2011; Yung et al., 2019). Similarly, endothelial cells influence different aspects of pancreatic tissue formation (Heymans et al., 2019; Jacquemin et al., 2006; Lammert et al., 2001; Lammert et al., 2003; Magenheim et al., 2011; Pierreux et al., 2010).

**Figure 1.**
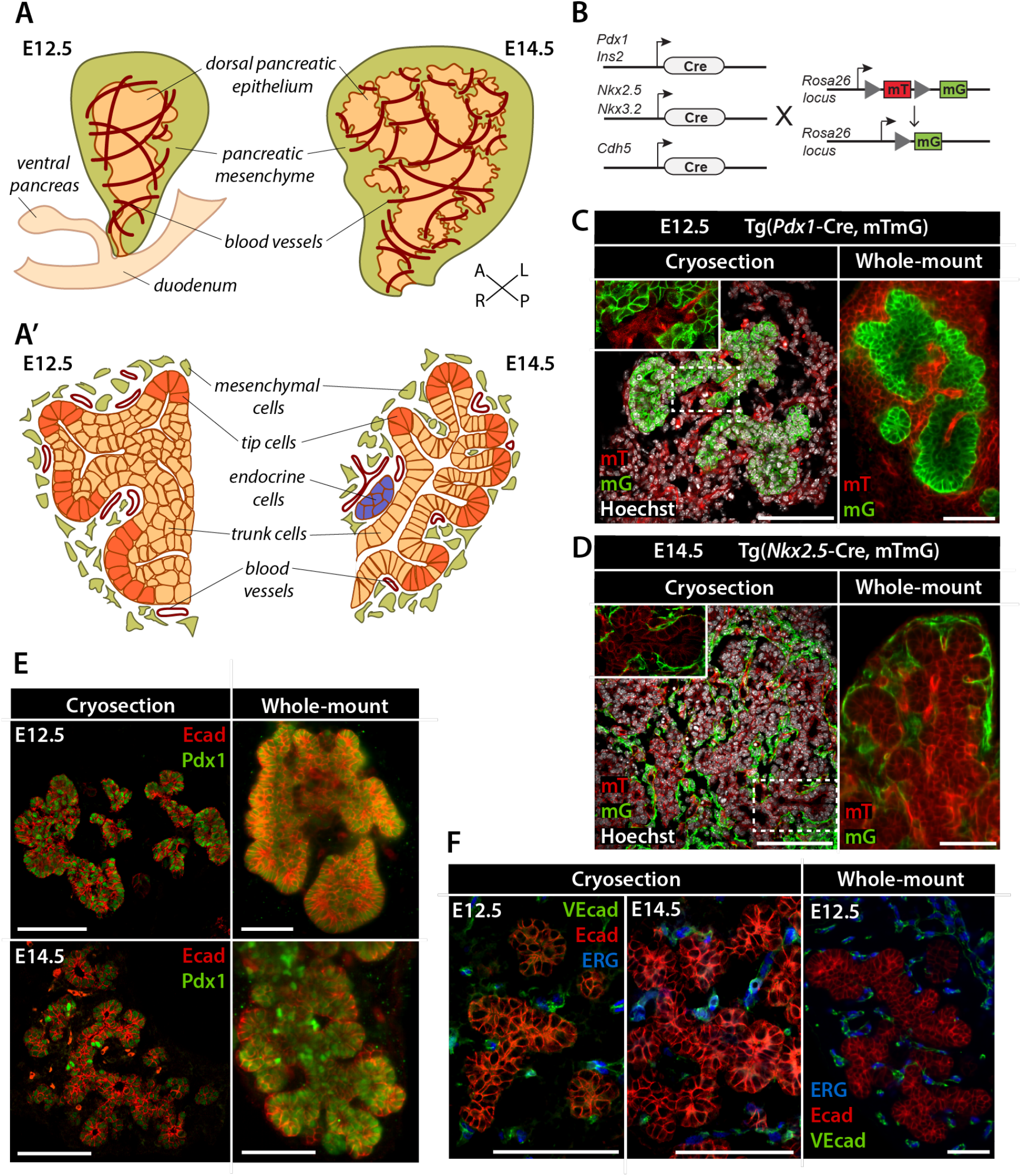
Analysis of pancreatic morphogenesis using confocal microscope and LSFM. **(A, A’)** Schematic representations of the pancreatic epithelium, spleno-pancreatic mesenchyme, and developing vasculature in E12.5 and E14.5 mouse embryos in 3D (A) or transverse section (A’). Depiction of tissue architecture in the transverse section highlights the segregation of the pancreatic epithelium into domains of “tip” and “trunk” cells from E12.5 onwards. A, anterior; P, posterior; L, left; R, right. **(B)** Genetic strategy for labeling distinct mouse pancreatic, endothelial, or mesenchymal cell populations. When not recombined, the transgenic (Tg) reporter construct mTmG results in the ubiquitous expression of a membrane-targeted tdTomato protein (mT). Cre-mediated recombination results in the excision of the mT expression cassette and expression of the membrane-targeted GFP (mG) in a tissue-specific fashion (Muzumdar et al., 2007). **(C, D)** Immunofluorescence (IF) detection of mG (green) and mT (red) in pancreatic tissue of E12.5 *Tg*(*Pdx1-Cre,* mTmG) embryos (C) or E14.5 *Tg*(*Nkx2.5-Cre,* mTmG) embryos (D). Pancreata analyzed as tissue sections by confocal microscopy (left panels) or as whole-mount preparations by LSFM (right panels) are shown side-by-side. In *Pdx1-Cre* Tg embryos (C), mG marks pancreatic epithelial cells. In *Nkx2.5-Cre* embryos (D), mG labels a subpopulation of mesenchymal cells in the spleno-pancreatic mesenchyme. Hoechst dye (grey) was used as nuclear counterstain on tissue sections. **(E, F)** Representative images of IF labelling of E12.5 and E14.5 pancreatic tissue. IF for E-cadherin (Ecad) (red, E and F) and Pdx1 (green, E) marks the pancreatic epithelium, while IF for ETS-related gene (ERG) (blue, F) and Vascular Endothelial cadherin (VEcad) (green, F) identifies endothelial cells surrounding the developing pancreas. Scale bars, 100 μm.

Temporal aspects of pancreas development have been elucidated in great detail, facilitated by time-resolved studies in various vertebrate models, including zebrafish, frog, chick, and mouse (Gittes, 2009; Larsen and Grapin-Botton, 2017; Zorn and Wells, 2009). These studies documented how and when early pancreatic progenitors commit to more specialized cell populations and how these processes can be recreated for *in vitro* differentiation and culture of pancreatic cell types. By contrast, spatial organization of the progenitor populations within the pancreas remains incompletely understood, because its embryonic development has been mainly studied in 2D on histological sections. Recent advances in microscopy techniques as well as optimized methods for clarification of whole tissue specimen have provided us with tools to study the pancreas in 3D at an unprecedented level of detail in the developing embryo (Chung et al., 2013; de Medeiros et al., 2016; Khairy and Keller, 2011; Richardson and Lichtman, 2015; Susaki et al., 2014; Wan et al., 2019). In particular, light-sheet microscopy has proven ideal for studying whole tissue samples in 3D as it combines excellent optical sectioning capabilities with fast image acquisition speed and reduced photoinduced damage to the tissue (de Medeiros et al., 2016; Khairy and Keller, 2011; Swoger et al., 2014; Wan et al., 2019).

Here, we applied light-sheet fluorescence microscopy (LSFM) to visualize the developing pancreas in 3D and to map tissue interactions at key time points during organ development. To this aim, we established protocols for tissue clarification and whole-mount immunofluorescence (WMIF) labelling of pancreatic tissue. We selected and validated transgenic mouse models and antibodies to visualize specific pancreatic epithelial, mesenchymal and endothelial cell populations and optimized light-sheet microscopy imaging protocols. In addition, we defined computational solutions for image analysis and quantification that enable detailed analysis of tissue composition and cell-cell interactions. Lastly, we combined our data in an open source online repository (referred to as Pancreas Embryonic Cell Atlas) to serve the scientific community by enabling further investigation of pancreas organogenesis and development of bioengineering solutions.

## Results

### Establishing the experimental tools to analyze pancreatic morphogenesis using light-sheet microscopy

Despite significant advancements in light-sheet microscopy, protocols for WMIF labelling, tissue clarification and imaging have not yet been established for the analysis of the developing pancreas and neighboring mesenchymal and endothelial tissues. Therefore, we started by testing antibodies and mouse reporter transgenic (Tg) strains for their suitability for 3D light-sheet microscopy of pancreatic tissue (Figs 1,2). Throughout the study, we compared data obtained by standard immunofluorescence labelling (IF) and confocal microscopy on pancreatic tissue sections with light-sheet microscopy images obtained from pancreata WMIF labelled for endogenous proteins or expressing fluorescent reporters.

**Figure 2.**
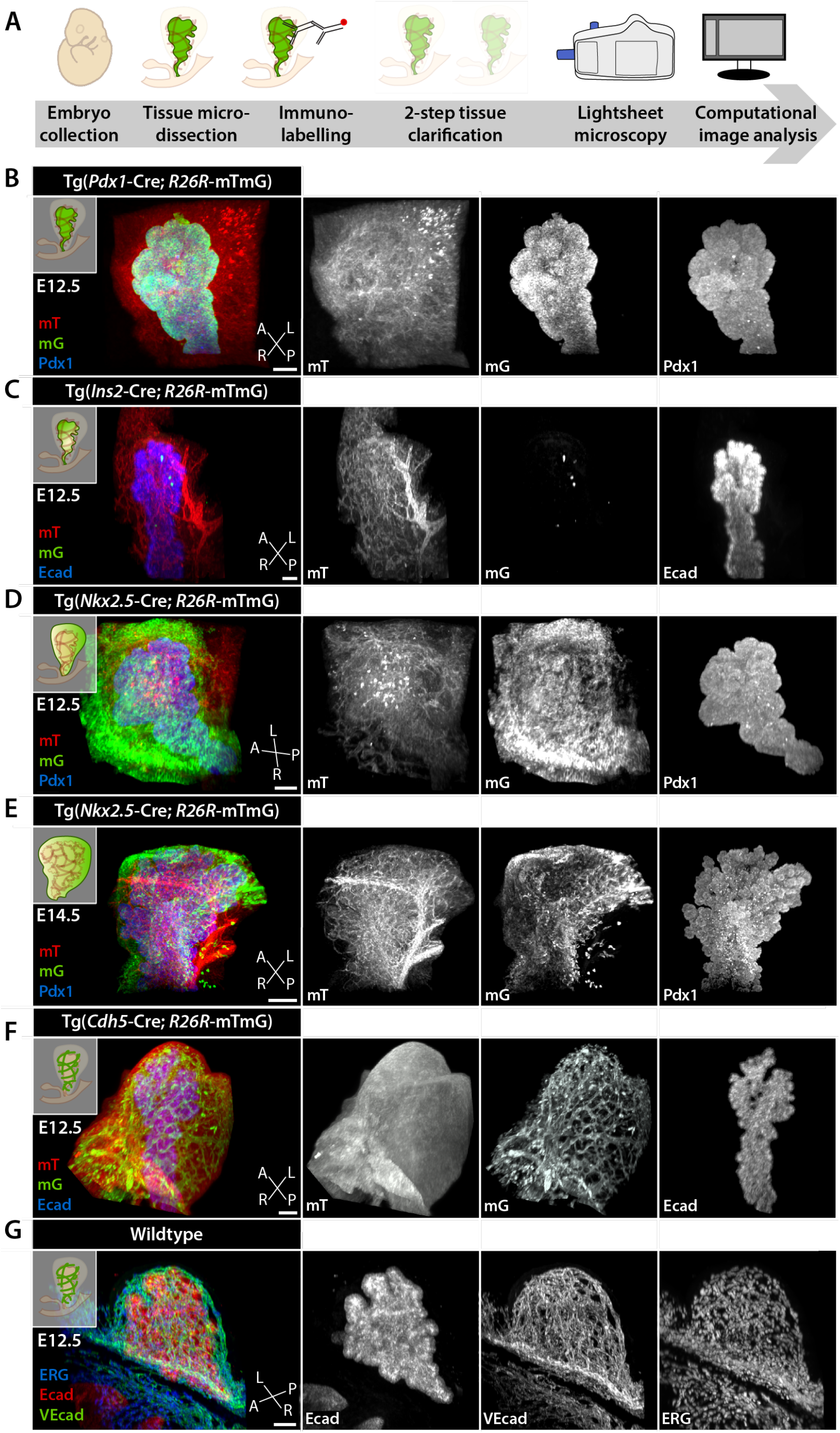
3D visualization of tissue interactions during pancreatic development using LSFM. **(A)** Schematic representation of the experimental setup for collection, visualization, and imaging of pancreatic tissue from mouse embryos using LSFM. **(B-G)** Representative LSFM 3D images of pancreatic tissue from transgenic (Tg) (B-F) or wild-type (G) mouse embryos at E12.5 (B-D, F, G) or E14.5 (E). Lineage-specific Cre induced mG expression in the indicated specific cell types. In *Pdx1-Cre* Tg embryos (B), mG (green) labels the entire pancreatic epithelium, while surrounding cells are mT^+^. In *Ins2-Cre* Tg embryos (C), mG marks insulin-expressing cells. In *Nkx2.5-Cre* Tg embryos (D and E), mG labels a sub-set of mesenchymal cells in the spleno-pancreatic mesenchyme. In *Cdh5-Cre* TG embryos (F), mG marks endothelial cells. In addition to detection of mG and mT, samples were stained for Pdx1 (B, D, E, blue) or Ecad (C, F, blue; G, red) to visualize pancreatic epithelial cells, for ERG (G, blue) and VEcad (G, green) to detect endothelial cells. A, anterior; P, posterior; L, left; R, right. Scale bars, 100 μm.

To spatially mark specific cell populations in the mouse embryonic pancreas, we used different Cre-driver lines to induce the expression of fluorescent reporter genes in epithelial, mesenchymal, or endothelial progenitor cell populations (Fig. 1B-D, Fig. 2). Reporter genes typically encode fluorescent proteins that can be visualized by detection of native fluorescence or immunolabelling with antibodies (Kretzschmar and Watt, 2012). For light-sheet microscopy and image segmentation, we chose the Tg(mTmG) dual fluorescent Cre reporter line, which is particularly suited for the visualization of cell-cell boundaries due to the membrane localization of the two reporter proteins (Kretzschmar and Watt, 2012; Muzumdar et al., 2007; Snyder et al., 2013). The membrane-localized tdTomato (mT) is constitutively expressed in all cells of the mouse embryo; after Cre-mediated recombination, the expression of membrane-localized EGFP (mG) replaces the mT cassette in all *Cre* expressing cells (Fig. 1B). We combined the Tg(mTmG) reporter line with five lineage-specific Cre Tg lines to label distinct cell types in the embryonic pancreas by the expression of mG (Fig. 1B). Specifically, we used the *Tg(Pdx1-Cre)* line to mark all cells of the pancreatic epithelium (Hingorani et al., 2003), the *Tg(Ins2-Cre)* line to mark insulin-expressing endocrine cells (Herrera et al., 1998), the *Tg(Cdh5-Cre)* line to mark endothelial cells (Chen et al., 2009), the *Tg(Nkx3.2-Cre)* and *Tg(Nkx2.5-Cre)* lines to label the entire spleno-pancreatic mesenchyme (Landsman et al., 2011; Verzi et al., 2009) and a subset of the mesenchymal cells surrounding the dorsal pancreas (Cozzitorto et al., 2020; Stanley et al., 2002), respectively.

Temporally, we focused our studies on the embryonic stages E12.5 and E14.5 (Fig. 1A), key time points in mouse pancreatic morphogenesis and differentiation (Larsen and Grapin-Botton, 2017; Pan and Wright, 2011; Shih et al., 2013; Zhou et al., 2007). At E12.5, the pancreatic epithelium converts from a relatively smooth, multilayered epithelial bud into a highly branched epithelial monolayer (Fig. 1A’). At E14.5, the pancreatic epithelium further differentiates, while pancreatic progenitors within the branches undergo proximo-distal patterning to become increasingly lineage restricted (Fig. 1A’).

First, we optimized currently available protocols for the preparation of mouse embryonic samples for LSFM to the pancreas (see also Methods section). For instance, to improve antibody penetration during WMIF labelling of embryonic pancreata, we extended the duration of the incubation steps with primary and secondary antibodies (Susaki et al., 2014). Moreover, to achieve complete tissue clarification, we modified the original CUBIC-based clarification protocols and the subsequent imaging setup (Susaki et al., 2014). Specifically, we subjected the WMIF-labelled pancreatic tissue to a prolonged two-step clarification procedure using CUBIC1 and CUBIC2 reagents, which resulted in optimal image quality (see Methods section for details).

To assess the image quality and resolution of our new protocols, LSFM of whole-mount pancreatic tissue from E12.5 *Tg*(*Pdx1-Cre;* mTmG) and E14.5 *Tg*(*Nkx2.5-Cre;* mTmG) embryos was compared to standard IF labelling and confocal microscopy on pancreatic tissue sections (Fig. 1C,D). 3D LSFM scan of E12.5 pancreas displayed primary branch structures composed of Pdx1^+^/mG-labelled epithelial cells surrounded by mesenchymal and endothelial tissues (mT) (Fig. 1C, right panel) with comparable resolution to 2D immunolabelled sections (Fig. 1C, left panel). Similarly, at E14.5 the pancreatic epithelial cells (mT) as well as Nkx2.5-Cre labeled mesenchymal cells (mG) were clearly visible using both imaging methods (Fig. 1D). Overall, these results indicate that our LSFM approach enables high-resolution imaging, comparable to conventional confocal microscopy approaches in 2D, but with the important advantage of preserving 3D tissue-architecture and avoiding technical artefacts due to tissue freezing, embedding and/or sectioning.

Next, we explored the potential of combining the detection of lineage-specific fluorescent reporters with IF for endogenous proteins in whole-mount to visualize specific cell populations. Antibodies against surface markers of epithelial [E-cadherin (Ecad)] and endothelial cells [VE-cadherin (VEcad)], as well as nuclear pancreatic [Pancreatic and duodenal homeobox 1 (Pdx1)] and endothelial [ETS-related gene (Erg)] transcription factors were tested in parallel in conventional confocal microscopy and LSFM imaging approaches. Both approaches showed consistent results enabling visualization of endogenous proteins in addition to the mG and mT labelled cells (Fig. 1E,F).

### 3D rendering of the developing pancreas enables qualitative assessment of tissue architecture

The main morphogenetic events underlying primary and secondary branch formation in the pancreas occur between E12.5-E14.5 in the mouse (Larsen and Grapin-Botton, 2017; Pan and Wright, 2011; Shih et al., 2013). So far, more attention has been paid to the intrinsic epithelial cell organization into defined branch shape and structure (Bankaitis et al., 2015; Hick et al., 2009; Kesavan et al., 2014; Kesavan et al., 2009; Petzold et al., 2013; Puri and Hebrok, 2007; Sznurkowska et al., 2018; Villasenor et al., 2010) than to the spatial organization of the different cell types, which surround the pancreatic branches. To fill this gap and capture the 3D structure of the whole organ, we acquired images of E12.5 and E14.5 whole-mount pancreata (Fig. 2). After image acquisition, the data were processed using ZEN and Imaris software to generate 3D representations of the acquired images (Fig. 2B-G). We generated a set of high-resolution images from E12.5 and E14.5 pancreata, in which epithelial and mesenchymal (Fig. 2B-E) or epithelial and endothelial cells (Fig. 2F,G) were simultaneously marked in the same embryo. This dataset provided us with suitable material to further analyze the interactions between the pancreatic epithelium and surrounding tissues.

Using the ‘Surface’ and ‘Spots’ components in Imaris, we reconstructed the pancreatic epithelium, mesenchyme and its vascular network in 3D and performed different analyses on the obtained 3D surface renderings, such as quantification of blood vessel diameter and distribution of endothelial/mesenchymal cells around the pancreatic epithelium (Figs 3,4). When comparing 3D surface renderings of the epithelium at E12.5 and E14.5, we observed an increase in branching complexity with time that was accompanied by a more densely packed blood vessels network wrapping the epithelium (Fig. 3A,B) as well as closer contacts of the Nkx2.5-Cre^+^ mesenchymal cells with the epithelium (Fig. 3C,D). Moreover, 3D surface renderings of the Nkx2.5-descendant mesenchymal population showed its preferential distribution along the left axis of the dorsal pancreatic epithelium (Fig. 3C,D), as previously reported by standard confocal image analysis (Cozzitorto et al., 2020).

**Figure 3.**
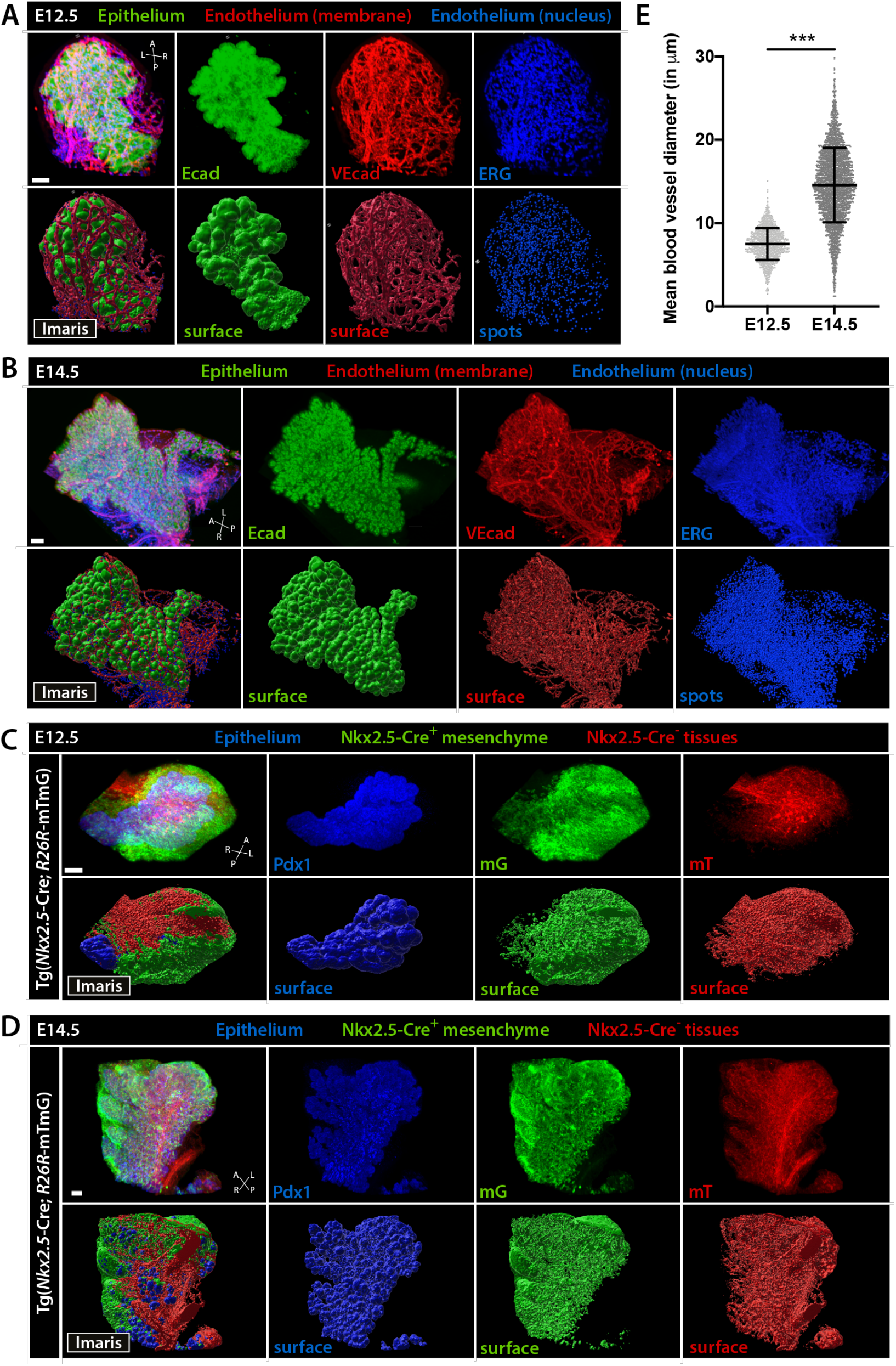
3D rendering of pancreatic tissue. **(A-D)** Representative LSFM 3D images (top panels) and Imaris surface renderings (bottom panels) of the developing pancreas from wild-type (A, B) or Tg(*Nkx2.5*-Cre; mTmG) (C, D) embryos at E12.5 and E14.5. WMIF for Ecad (A, B, green) or Pdx1 (C, D, blue) labels the pancreatic epithelium and for VEcad (red; A, B) or ERG (blue; C, D) marks the endothelium. mG (green; C, D) and mT (red; C, D) in Tg(*Nkx2.5*-Cre; mTmG) embryos mark the Nkx2.5-Cre^+^ mesenchyme and Nkx2.5-Cre^-^ tissues, respectively. 3D images and surface renderings are shown as merged (leftmost panel) and individual channels (right panels). A, anterior; P, posterior; L, left; R, right. Scale bars, 100 μm. **(E)** Quantification of the mean diameters of blood vessels in close proximity (<15 μm) to the pancreatic epithelium at E12.5 and E14.5. Mann-Whitney test, p-value < 0.001.

**Figure 4.**
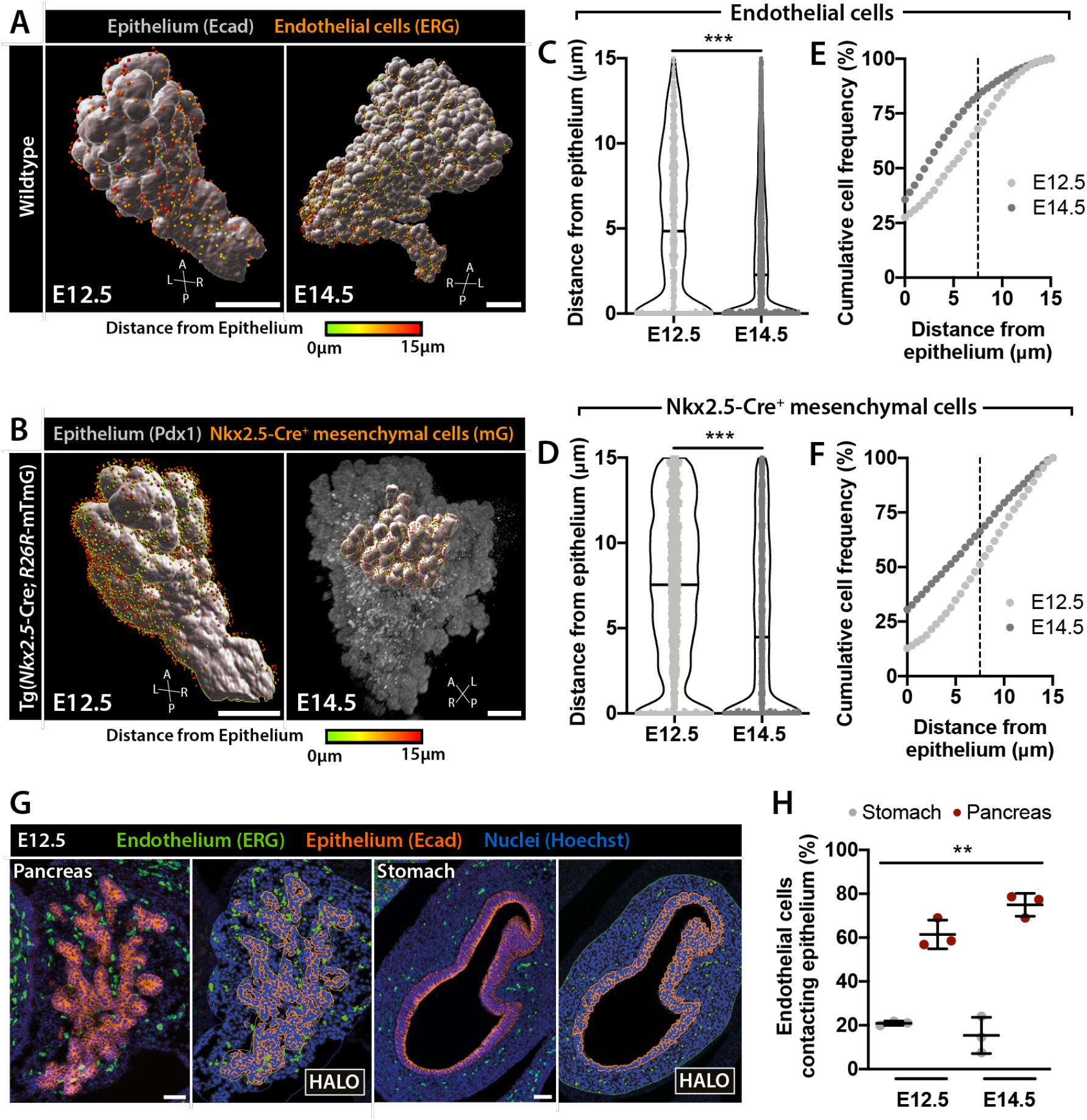
Quantitative analysis of endothelial and mesenchymal cell distribution around the pancreatic epithelium. **(A, B)** 3D rendering of LSFM scans of E12.5 and E14.5 pancreas and surrounding tissues labelled with indicated antibodies. Ecad^+^ (A) or Pdx1^+^ (B) pancreatic epithelial cells were rendered as surfaces (grey). ERG^+^ endothelial (spots, A) or Nkx2.5-Cre^+^ mesenchymal (spots, B) cells were identified using ‘Spots’ function in Imaris and color-coded based on their distance from the epithelium. Only endothelial or mesenchymal cells within a 15 μm distance from the epithelium were considered in the analysis. A, anterior; P, posterior; L, left; R, right. Scale bars, 200 μm. **(C, D)** Violin plots showing the distribution of endothelial (C) or Nkx2.5-Cre^+^ mesenchymal cells (D) around the pancreatic epithelium at E12.5 [n=799 cells (C); n=3350 cells (D)] and E14.5 [n=7817 cells (C); n=3359 cells (D)]. Mann-Whitney test, p-value <0.001 (C and D). **(E, F)** Analysis of endothelial (E) or Nkx2.5-Cre^+^ mesenchymal cell (F) distance from pancreatic epithelium presented as cumulative frequency distribution. The percentage (%) of endothelial (E) or Nkx2.5-Cre^+^ mesenchymal (F) cell population is plotted against the distance from the epithelium. Dashed vertical line indicates the 7.5 μm boundary, which corresponds approximately to the thickness of the first cell layer in direct contact with the epithelium. **(G)** Representative images of Ecad (orange) and ERG (green) IF labelling in E12.5 pancreas and stomach sections. Hoechst (blue) was used as nuclear counterstain. Scale bar, 50 μm. **(H)** Scatter plot showing quantification of endothelial cell distance from the pancreas or stomach epithelium at E12.5 (n=3) and E14.5 (n=3). Quantification was performed using HALO software. Data represents the number of endothelial cells localized within the first cell layer from the epithelium as % of the total number of endothelial cells within a distance < 15 μm from the pancreas or stomach. Kruskal-Wallis test, p-value=0.001.

### LSFM imaging enables quantitative analysis of tissue organization during development

LSFM images are a source of qualitative but also quantitative information that might complement 2D observations, inform on pancreas organogenesis, or generate image-driven hypotheses. For instance, 3D reconstruction of the VEcad^+^ vascular network at E12.5 and E14.5 allowed us to measure the diameter of the blood vessels in close proximity to the pancreatic bud (up to 15μm from the epithelial surface). We found a two-fold increase in the average diameter of blood vessels closely associated with the pancreatic epithelium between E12.5 and E14.5 (Fig. 3E). These findings support the notion that a perfused and mature vascular network is established in the pancreas between E12.5-E14.5, as previously suggested (Shah et al., 2011).

Additionally, we quantified the distribution of endothelial and Nkx2.5-Cre^+^ mesenchymal cells with respect to their distance from the epithelial surface and analysed its changes over time. 3D renderings of the pancreatic epithelium surface were based on Ecad (Fig. 4A) or Pdx1 (Fig. 4B) WMIF images, while the Imaris ‘Spots’ function enabled identification of endothelial or Nkx2.5-Cre^+^ mesenchymal cells based on ERG (Fig. 4A) and mG (Fig. 4B) expression, respectively. We limited our analysis to endothelial and mesenchymal cells within a range of 15 μm from the epithelial surface, as cells in close proximity are likely to influence morphogenetic events of the underlying epithelium through paracrine signalling or physical constraint. Comparison of the distribution of endothelial and Nkx2.5-Cre^+^ mesenchymal cells within the 15 μm radius highlighted a general trend toward increased density of both cell populations in proximity with the pancreatic epithelium as development progresses (Fig. 4C,D). By plotting the cumulative frequency distribution for endothelial and mesenchymal cells against their distance from the epithelium, we measured about 68% of endothelial and 51% of Nkx2.5-Cre^+^ mesenchymal cells present within the first cell layer (about 0-7.5μm) at E12.5 and 82% of endothelial and 66% of Nkx2.5-Cre^+^ mesenchymal cells at E14.5 (Fig. 4E,F).

To validate and further expand the analysis of endothelial cell distribution in relation to the pancreatic epithelium, we analysed serial IF labelled tissue sections and segmented epithelial and endothelial cells using the HALO software (Fig. 4G). Quantification of the percentage of endothelial cells within the first and second cell layer around the pancreatic epithelium documented about 61% of endothelial cells in direct contact with the epithelium at E12.5 and 75% at E14.5 (Fig. 4H), thereby corroborating the results obtained from 3D images (Fig. 4C,E). Interestingly, equivalent analysis of the vasculature around the stomach epithelium revealed significantly lower proportions of endothelial cells contacting the gastric epithelium at E12.5 and E14.5 (21% and 15%, respectively). These results suggested a tissue-specific vascularization pattern, whereby the pancreatic epithelium might require a closer interaction with a dense endothelial network to successfully undergo the complex morphological changes that occur during these developmental stages.

Overall, these analyses underscore previously underappreciated spatio-temporal changes in the composition and organization of the immediate pancreatic microenvironment and warrant further investigations.

### 3D image analyses allow precise quantification of cell type abundance

Numerous studies previously highlighted the importance of a finely tuned balance between epithelial, endothelial, and mesenchymal cell types to allow pancreatic growth and morphogenesis. For instance, the presence of an excessive number of endothelial cells has been shown to impair pancreatic growth and limit branching morphogenesis (Magenheim et al., 2011; Sand et al., 2011) as well as to promote endocrine islets hyperplasia and reduce acinar differentiation (Lammert et al., 2001; Pierreux et al., 2010). By contrast, depletion of mesenchymal cells leads to severe pancreatic hypoplasia (Landsman et al., 2011). Therefore, establishing the relative abundance of epithelial, endothelial, and mesenchymal cell populations during pancreatic development is critical for the ongoing efforts to bioengineer pancreatic tissues through co-culture or 3D bioprinting of heterologous cell populations.

To measure the relative abundance of the three cell types in the developing pancreas, we used 3D LSFM scans of E12.5 and E14.5 pancreata labelled for Ecad to mark epithelial cells and ERG to mark endothelial cells (Fig. 5A). DRAQ5 was used as nuclear counterstain. Using the ‘Spots’ function in Imaris, we quantified the number of epithelial (Ecad^+^/ERG^-^/DRAQ5^+^), endothelial (Ecad^-^/ERG^+^/DRAQ5^+^), and mesenchymal (Ecad^-^/ERG^-^/DRAQ5^+^) cells within a 15 μm distance from the epithelial surface, as in the cell distribution analysis (Fig. 4). We found that the proportions of epithelial (~66%), endothelial (~7%), and mesenchymal (~27%) cells remain stable within this extended pancreatic region between E12.5 and E14.5 (Fig. 5C). It should be noted that in this analysis a mesenchymal cell identity was assigned based on the absence of epithelial and endothelial markers (Fig. 5), which does not exclude the presence of other cell types, such as neuronal, lymphatic, or immune cells, in this fraction. However, these cell populations are rare within the pancreatic tissue at E12.5 and E14.5 (Burris and Hebrok, 2007; Cozzitorto et al., 2020) (Fig. S2), especially within the 15 μm radius from the epithelium, being therefore negligible in the calculation of relative cell abundance.

**Figure 5.**
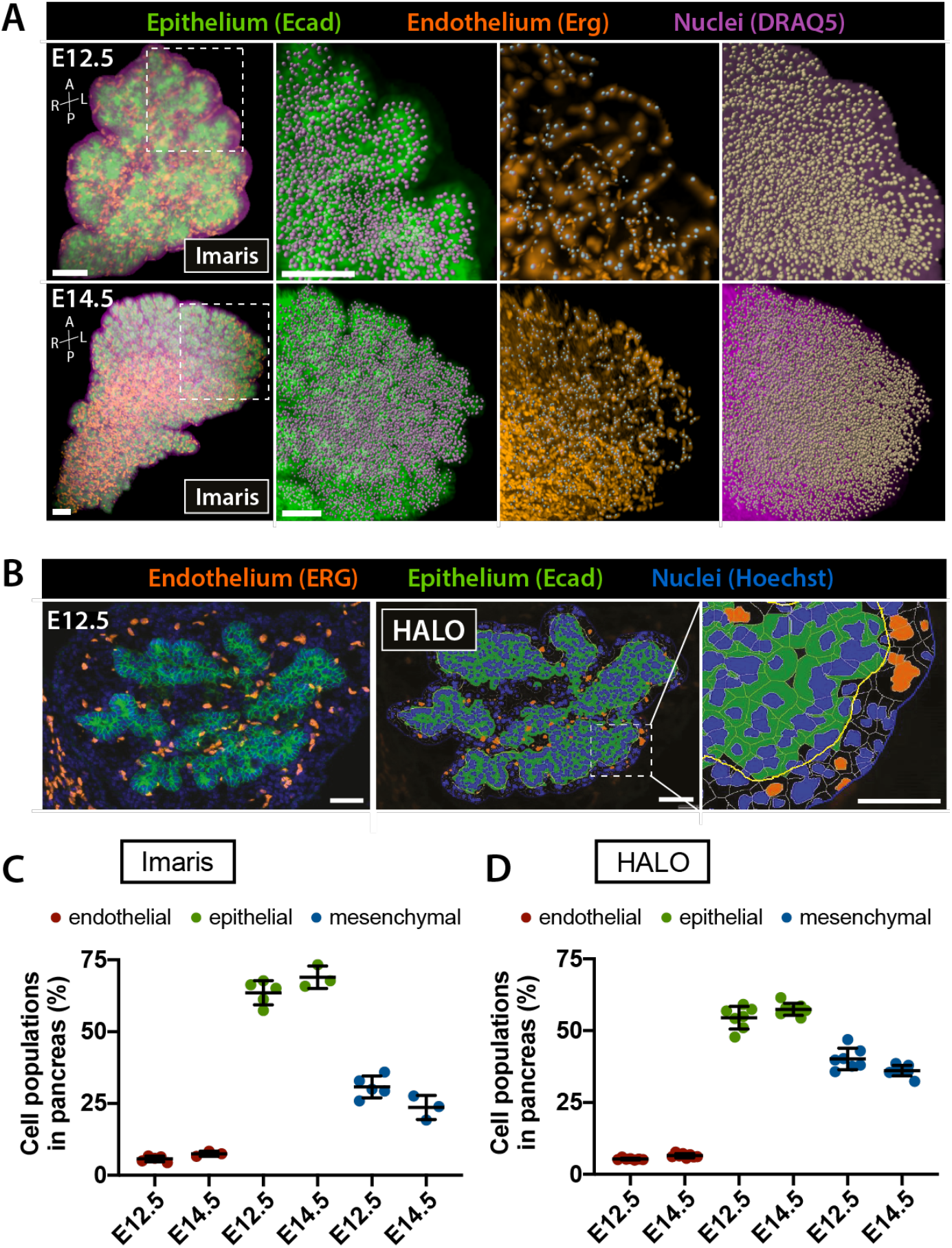
Tissue segmentation of murine embryonic pancreas. **(A)** 3D representation of LSFM scans of E12.5 and E14.5 pancreas and surrounding tissues labelled with antibodies against Ecad (epithelial cells, green) and ERG (endothelial cells, orange). DRAQ5 was used as nuclear counterstain. ‘Spot detection’ function in Imaris was used to identify all cells (based on DRAQ5), individual epithelial (Ecad^+^) or endothelial cells (ERG^+^) in the indicated boxed regions. Mesenchymal cells were identified as DRAQ5^+^/Ecad^-^/ERG^-^. Analysis was restricted to surrounding tissue within 15 μm distance from the epithelial surface. Right panels show higher magnification single-channel images of the boxed region. Scale bars, 100 μm. **(B)** Representative image of Ecad and ERG IF labelling of E12.5 pancreatic tissue sections (left panel). Hoechst was used as nuclear counterstain. Epithelial, endothelial, and mesenchymal cells were segmented using HALO software (middle panel). Right panel shows higher magnification of the boxed region. Mesenchymal cells were identified as Ecad^-^/ERG^-^/Hoechst^+^. Scale bars, 50 μm; 20 μm (magnification panel). **(C, D)** Scatter plots of relative abundance of endothelial, epithelial, and mesenchymal cells within a 15 μm radius of pancreatic tissue, shown as % of total cell numbers. Analysis of cell type fractions in (C) was performed using Imaris software on LSFM of whole-mount samples (A), while analysis in (D) was performed using HALO software on confocal images of IF sections (B). No differences in cell fractions were detected between embryonic stages (E12.5 *vs* E14.5) or analysis pipelines (Imaris *vs* HALO). Kruskal-Wallis test, followed by Dunn’s multiple comparisons test.

Next, we corroborated these results by assessing the relative abundance of the three cell populations on IF labelled sections spanning the entire tissue and segmenting epithelial, endothelial, and mesenchymal cells using the HALO software (Fig. 5B). Analogous to our previous analysis, we restricted our analysis to tissues within a 15 μm range from the epithelium. We used nuclear counterstain to mark all cells within the region of interest (ROI) in combination with antibodies against Ecad to mark epithelial and ERG to mark endothelial cells. The values in the proportions of epithelial (~56%), endothelial (~6%), and mesenchymal (~38%) cells remained constant between E12.5 and E14.5 (Fig. 5D). Notably, we observed only minor discrepancies in the percentages of epithelial (~66% *vs* ~56%) and mesenchymal cells (~27% *vs* ~38%) when comparing results obtained from 3D LSFM scans and 2D confocal images. This is likely due to differences in cell segmentation between the Imaris and HALO software used for the image analysis.

In conclusion, we used 3D and 2D image analysis pipelines to assess the relative abundance of the three main cell types in the pancreatic tissue during development. Interestingly, our findings indicate that despite profound morphological changes in tissue architecture between E12.5 and E14.5, the relative cell type composition in the pancreatic tissue does not change. Hence, an optimal ratio of epithelial (~60%) to endothelial (~6%) to mesenchymal (~34%) cells might be required to support pancreatic development.

### The Pancreas Embryonic Cell Atlas as an open access data repository

All 3D LSFM images visualizing the three main cell types (epithelium, endothelium and mesenchyme) of the murine embryonic pancreas have been annotated and deposited on the open-source database openBIS (open Biology Information System) (https://openbis.ch/) to start building a Pancreas Embryonic Cell Atlas (Table S1). We built a customized data management platform and all image data have been made available on the data management platform openBIS (https://openbis-data-repo.ethz.ch/openbis/). So far, 51 LSFM scans have been made publicly available in our data repository openBIS in the following formats: tiff (raw microscopy data), ims (proprietary format of Imaris containing microscopy images and surfaces with tissue shapes), wrl (tissue shapes exported separately), png (overview of the sample). Each image collection comprises a context-specific group of images and annotations, including developmental stage, mouse genotype, mouse background, segmented tissue, antibodies and conjugated fluorescent dyes (450 nm/488 nm/594 nm/647 nm), laboratory of origin, site of imaging and comments. Modalities on how to access the data are in the Methods Section and Fig. S1.

## Discussion

Spatial information on the interaction of pancreatic progenitor cell types with surrounding tissues is essential for the full comprehension of basic concepts governing pancreas development *in vivo* but also for devising strategies for the *in vitro* generation of transplantable pancreatic tissue for regenerative therapies. To date, spatial information on pancreas development has mostly relied on 2D histological sections or *in silico* reconstructions (Bankaitis et al., 2015; Bhushan et al., 2001; Heymans et al., 2019; Kesavan et al., 2014; Kesavan et al., 2009; Mamidi et al., 2018; Villasenor et al., 2010). Previous studies of pancreatic tissue in whole-mount have provided excellent information on the gross morphology of the epithelium (Fowler et al., 2018; Kesavan et al., 2009; Petzold et al., 2013; Sznurkowska et al., 2018; Villasenor et al., 2010) but limited information on contacts and interactions with neighboring tissues. Nowadays, light-sheet microscopy has revolutionized the study of complex organ anatomy enabling to visualize the architecture of developing or adult tissues in 3D (de Medeiros et al., 2016; Khairy and Keller, 2011; Roostalu et al., 2020; Swoger et al., 2014; Wan et al., 2019). Here, we have established tools and protocols for WMIF labelling, tissue clarification, and light-sheet imaging of the developing pancreas and exemplified pipelines to utilize these data to the study of pancreatic organ development in the mouse.

Currently, heterologous tissue interactions, such as epithelial-mesenchymal or epithelial-endothelial crosstalk during pancreas development, has gained a lot of attention because of their essential role in pancreatic cell differentiation (Attali et al., 2007; Azizoglu and Cleaver, 2016; Cozzitorto et al., 2020; Heymans et al., 2019; Landsman et al., 2011; Pierreux et al., 2010; Sakhneny et al., 2019; Seymour and Serup, 2019; Yung et al., 2019). As a proof-of-concept, here we exemplified how to extract information on the 3D tissue architecture as well as the relative position of epithelial, mesenchymal, and endothelial cells from LSFM images of the developing pancreas. These data provide us with novel ways to visualize the intimate interaction between epithelial cells and Nkx2.5^+^ mesenchymal cells, which contribute to the proper development of the endocrine pancreas (Cozzitorto et al., 2020). Furthermore, we applied our analysis pipelines to gain insights into the cell ratio between epithelial, mesenchymal, and endothelial cells at key time points during pancreas development. This information will be instrumental in assisting current efforts to improve multi-tissue bioengineering approaches aimed at advancing the physiological complexity in *in vitro* pancreatic culture systems.

We have provided access to our acquired dataset through the Pancreas Embryonic Cell Atlas, an open source data repository (Fig. S1). So far, 3D LSFM-based analyses of the mouse embryonic pancreas together with its neighboring mesenchymal and endothelial tissues have not been reported. Thus, the Pancreas Embryonic Cell Atlas will provide the scientific community with a valuable source of data for future investigation. Similar to other open source data repositories, such as the human ‘Pancreatlas’ (https://www.pancreatlas.org/) (Saunders et al., 2020), we envision the Pancreas Embryonic Cell Atlas as a dynamic platform that will evolve with continued input from us and others. Foremost, the collection of raw data deposited in the Pancreas Embryonic Cell Atlas will help the scientific community to develop and test novel hypotheses on tissue interactions that guide pancreatic differentiation and morphogenesis. Also, we provide datasets of already rendered surfaces, as exemplified in Fig. 3, that can be viewed using the free Imaris Viewer software and do not require access to commercial software or high-performance computing.

Another exciting application we envision for our dataset is its value for approaches aimed at bioprinting pancreatic tissues. Bioprinting is an innovative technology that enables the assembly of multiple biological components (cells and biomaterials) in a precise ratio and controllable anatomical geometry, and with high-throughput capability and reproducibility (Shafiee and Atala, 2017). Conceptually, the information on pancreatic tissue architecture gained in this study provides the blueprint for 3D bioprinting of the diverse cell types that comprise this multi-tissue pancreatic organ rudiment. In particular, future studies will use the image data files deposited in the Pancreas Embryonic Cell Atlas to generate CAD files for developing bioprinting applications to generate pancreatic tissue or functional islet-like structures for cell therapy to treat diabetes or for pancreatic disease modeling. Finally, we envision that Pancreas Embryonic Cell Atlas image collections will identify collaborations also outside the pancreas field and lay the foundation of interdisciplinary work that integrates cellular-resolution data within the context of whole-organ architecture.

## Methods

### Animal experimentation

The following transgenic mouse lines were used: Tg(*PdX1*-Cre) (Hingorani et al., 2003), *Tg(Nkx3.2-Cre)* (Verzi et al., 2009), Tg(*Nkx2.5*-Cre) (Stanley et al., 2002), Tg(*Ins2*-Cre) (Herrera et al., 1998), Tg(*Cdh5*-Cre) (Chen et al., 2009), *Tg(R26R-* mTmG) (Muzumdar et al., 2007). All mouse strains were on a C57BL/6 genetic background and kept under standard housing conditions. All procedures relating to animal care and treatment conformed to the Institutional Animal Care and Research Advisory Committee and local authorities (PPL PP6073640, Home Office, UK; University Animal Welfare Committee UCLouvain 2016/UCL/MD/005 and 2020/UCL/MD/011; Veterinary Office of Canton Basel-Stadt, Switzerland; Tel Aviv University Committee on Animal Research). For timed mating, male and female mice were placed into a breeding cage overnight and plug check was performed daily. The presence of a vaginal plug in the morning was noted as embryonic day (E) 0.5. Embryos were collected at E12.5 and E14.5 and dissected under a stereomicroscope.

### Immunofluorescence labelling of mouse tissue sections

After dissection, embryos were fixed in 4 % paraformaldehyde (PFA) in phosphate buffered saline (PBS) overnight at 4°C. For generation of paraffin sections fixed embryonic tissue was embedded in paraffin using a Tissue-Tek VIP-6 (Sakura). For generation of cryosections fixed embryonic tissue was equilibrated overnight in 20% sucrose solution and embedded in O.C.T. compound (Tissue-Tek, Sakura). Paraffin- and cryosections were cut at 7 μm and 10 μm thickness, respectively. Prior to immunolabelling, paraffin sections were deparaffinized with xylene for 10min and rehydrated. If required, antigen retrieval was performed using citrate buffer. Next, tissue sections were incubated in TSA (Perkin Elmer) blocking buffer for 1 hour (h) at room temperature followed by overnight treatment at 4° C with primary antibodies at the appropriate dilution (Table S2). Next, tissue sections were incubated for 1 h at room temperature with Hoechst 33342 nuclear counterstain at a concentration of 250 ng/ml and secondary antibodies at the appropriate dilution (Table S2). Slides were mounted with Dako fluorescent mounting medium and imaged on a Zeiss LSM 700 confocal microscope using a 25x or 40x oil immersion objective.

### Immunofluorescence labelling of embryonic tissue in whole-mount

Embryos were collected at E12.5 and E14.5 and the trunk region posterior to the forelimbs and anterior to the hindlimbs was dissected. The duodenum, stomach, dorsal and ventral pancreas as well as associated mesenchymal tissues were dissected and harvested as a continuous unit. Optionally, to simplify the image acquisition of the pancreas at E14.5, the stomach was removed. The dissected tissues were fixed in 4 % PFA for 1-2 h at room temperature and then extensively washed in 1X PBS. Subsequently, samples were placed in blocking solution [3 % donkey serum (DS), 0.1 % (at E12.5) or 0.5 % (at E14.5) Triton X-100 in 1X PBS] for 32 h at 4° C and afterwards incubated with primary antibodies in blocking solution at the appropriate dilution for 48 h at 4°C (see Table S2). After washes in freshly prepared 1X PBS with 0.1% Triton X-100 solution (at least 3 times, 5 min each, followed by an overnight washing step at 4°C), the samples were incubated with secondary antibodies appropriately diluted (see Table S2) together with Hoechst 33342 nuclear counterstain (250 ng/ml) in blocking solution for 32 h at 4° C. After extensive washing steps in 1X PBS with 0.1% Triton X-100 (as above), the samples were transferred into PBS and stored at 4° C until tissue clarification.

### Tissue clarification

Pancreatic tissue was cleared in freshly prepared CUBIC1 (25 % wt/vol urea, 25 % wt/vol N,N,N’,N’-tetrakis(2-hydroxypropyl) ethylenediamine, 15 % wt/vol Triton X-100, in dH_2_O) and CUBIC2 (50 % wt/vol sucrose, 25 % wt/vol urea, 10 % wt/vol 2,20,20’-nitrilotriethanol, 0.1 % vol/vol Triton X-100, in dH_2_O) solutions as previously described (Susaki et al., 2014).

The clarification protocol was adapted according to the mounting procedure used for light-sheet microscopy. If to be glued to a supportive holder, the sample was transferred into a glass vial containing 2 ml CUBIC1 solution and incubated for 3 weeks at room temperature on a rotational shaker in the dark. The tissue was then transferred into a new glass vial containing 2 ml of CUBIC2 solution, without carrying over any CUBIC1 solution, and incubated for 1 week at room temperature on a rotational shaker in the dark.

If to be embedded in agarose, the sample was immersed in CUBIC1 at 37°C for 3 days in plastic Petri dishes or 4-well plates on a rotational shaker, then washed in PBS for 1 day at 4° C and transferred in CUBIC2 at room temperature for 3 weeks with solution changed every 3-5 days. During the PBS washing step, samples were again incubated with Hoechst 33342 nuclear counterstain (250 ng/ml). Both CUBIC1 and CUBIC2 incubation steps were preceded by an intermediate 1:2 dilution step of the respective solution of 24 h duration.

### Sample mounting and light-sheet microscopy

Samples were glued to a 1 ml syringe supportive holder using all-purpose super glue. The syringe was then inserted into the syringe holder provided with the Zeiss Z1 light-sheet microscope. Specifically, the clarified tissue was retrieved from the CUBIC2 solution by removing as much solution as possible using either a pipette or paper wipes and glued to the tip of the syringe. To avoid interference with pancreatic tissue during imaging, the sample was glued on the stomach or duodenum. After 1 minute (min), the sample holder was inserted it into the microscope for image acquisition. Alternatively, for agarose mounting, the samples were embedded in a glass capillary, provided by Zeiss, filled with 2% low-melting point agarose solution. Afterwards, the solidified agarose cylinders containing the samples were immersed in CUBIC2 for 48 h to clear the agarose. Samples were then imaged with the Zeiss Z1 light sheet microscope using 20x acquisition lens and 10x illumination lenses.

### Image analysis and cell segmentation

Imaris (Bitplane Oxford Instruments, version 9.5.1) software packages, including Imaris XT, Imaris Filament Tracer, Imaris Vantage and Imaris Measurement Pro with IPSS and Statistics features, were used to segment the 3D images obtained from the light-sheet microscope. Volumes of the different cell populations in the developing pancreas were created using the ‘Surface creation’ module on the epithelial [E-cadherin (Ecad) or Pdx1 labelling], the endothelial [VE-cadherin (VEcad) labelling] and the Nkx2.5-Cre^+^ mesenchymal populations. Nuclei of endothelial and Nkx2.5-Cre^+^ mesenchymal cells were reconstructed using the ‘Spots creation’ module and used as the center of the cell, based on the endothelial ERG signal and the inverted signal of the Nkx2.5-Cre labeled mesenchymal cells (mG), respectively.

The mean vascular diameter was computed using the ‘Filament tracer’ extension on the VEcad labelling surrounding the epithelium bud within a distance of 15 μm. The mean diameter of each vascular segment between two junctions was extracted with the ‘Vantage’ Extension package. The distribution of endothelial and *NKxv2.5*-expressing mesenchymal cells in the vicinity of the pancreatic bud was performed using the reconstructed volume of the epithelial cells and the spots generated for the endothelial and Nkx2.5-Cre^+^ mesenchymal cells. Using the feature ‘Shortest distance’, the distance of each endothelial or Nkx2.5-Cre^+^ mesenchymal cells from the epithelial surface was extracted. Because of the complex morphology of the pancreas with its multiple branches, some endothelial and Nkx2.5^+^ cells appeared included in the epithelium volume. These cells were actually located in spaces between pancreatic branches and were considered at a distance of 0 μm, as they were in direct contact with epithelial cells.

To segment pancreatic tissue into single cells for cell type quantification, LSFM scans of pancreata labelled with antibodies against E-cadherin (Ecad) and ERG and DRAQ5 nuclear counterstain were analysed at defined ROI [300×300×300 μm (E12.5 pancreata) or 400×400×400 μm (E14.5 pancreata)]. Before proceeding with the segmentation, the Ecad membrane signal was subtracted from the DRAQ5 signal to obtain a sharper delineation of the nuclei. Additionally, the Ecad signal was blurred with a ‘Gaussian filter’ and inverted to obtain a filled cell signal with dark membranes. This multi-step image processing allowed us to identify three distinct types of ‘Spots’ in Imaris: one for epithelial cells based on the inverted Ecad signal; one for endothelial cells based on the ERG nuclear signal; one for all nuclei based on the DRAQ5 signal (identification settings for all were set to a XY diameter of 5 μm and a Z diameter of 7 μm). To restrain the segmentation analysis to the epithelial bud and tissues in its immediate vicinity, an epithelial surface was created, as described above. Then, a filtering step was applied to exclude all nuclei and endothelial spots found at a distance greater than 15 μm from the epithelial surface. Next, an additional filtering step was applied to remove undesirable spots, such as multiple spots found in larger cells. Finally, the whole-organ segmentation process was verified by extracting virtual 2D sections of the fluorescence signals and their respective spot signals and by applying manual correction where necessary.

The image analysis software HALO (Indica Labs, v3.0.311.317) was used to segment images from tissue sections of E12.5 and E14.5 pancreatic and gastric tissue IF-labelled with Ecad and ERG antibodies and Hoechst nuclear counterstain. First, the area of interest was defined by manually delineating the pancreatic epithelium as well as the surrounding tissues within a distance of approximately 15μm (about 2 cell layers) from the epithelium. The CytoNuclear FL v1.4 Algorithm was applied to segment the tissue into individual cells. This algorithm segments the nuclei based on Hoechst staining and further identifies distinct cell types based on the fluorochrome detected in the segmented area, *i.e*. Ecad marking epithelial cells and ERG marking endothelial cells. To quantify the distribution of endothelial cells around the pancreatic or gastric epithelium, the XY coordinates for endothelial and epithelial cells were extracted from the segmented images. Using R software, the number of endothelial cells in direct contact with the epithelium (0 μm-7.5 μm distance from the epithelium) and within a 15μm distance from the epithelium was computed. The relative abundance of epithelial, endothelial, and mesenchymal cells was determined by quantifying the number of each cell type on regularly spaced sections (every 30 μm) spanning the entire pancreatic tissue.

### Pancreas Embryonic Cell Atlas and Browsing Features

The images are stored on the data management platform openBIS (https://openbis.ch/). For repository access, first please contact the PAN3DP consortium (https://www.pan3dp-project.eu/ or) to be granted guest access to the platform and then visit (https://openbis-data-repo.ethz.ch/openbis/). The collections are accessible within the Iber Materials/Publications/Group Publications/Pancreas Atlas. Each collection corresponds to one developmental stage. Within each collection, the objects correspond to specific genotypes, *i.e*. transgenic mouse lines. Each object contains individual samples, *i.e*. the image datasets. For a schematic representation of data organisation in the repository, see Fig. S1. Within each sample, images in all different formats can be retrieved using advanced filter commands and downloaded separately or collectively. The data can be quickly accessed for viewing and download as zip folder. The information displayed in the ‘Results’ table can be adapted according to individual needs by using the ‘Columns’ drop-down menu.

To have the most informative overview on the 3D architecture of the pancreas, it is recommended to view the surfaces combined with the multi-channel microscope images in the Imaris proprietary format (.ims) using the free Imaris Viewer (https://imaris.oxinst.com/imaris-viewer). To perform further modification on the data, the commercial version of Bitplane Imaris software is required. The raw .tiff files can be opened with open-source software such as ImageJ or Fiji. The surface files in .wrl format can be opened using 3D mesh processing software, such as MeshLab.

## Competing interests

No competing interests declared.

## Acknowledgements

We acknowledge the support of the European Union’s Horizon 2020 research and innovation programme Pan3DP FET Open [grant Number 800981]. LG holds a fellowship from the Fonds pour la formation à la Recherche dans l’Industrie et l’Agriculture (FRIA, Belgium). We thank Steve Runser from the Iber lab. and Caterina Barillari from SIS, ETHZ for setting up the openBIS infrastructure for the pancreas atlas and helping with creating the public download link.

## Authors contributions

LG, AS and DW designed and performed experiments, analysed data and wrote the manuscript. MM, JFD, SE, LS, AS, LS performed experiments; HG, CL, LC contributed to segmentation and image analysis. LL, SL, FG are PIs of the Pan3DP consortium, contributed intellectually to the project and funding acquisition. DI, CP and FMS are PIs of the Pan3DP consortium, contributed intellectually to the project and funding acquisition, co-supervised the project; FMS finalized the manuscript.

---

**Table 1.** Proportion of endothelial, epithelial and mesenchymal cells in murine embryonic pancreas.

|  | **E12.5 (%)** |  |  | **E14.5 (%)** |  |  |
| --- | --- | --- | --- | --- | --- | --- |
|  | Endothelial | Epithelial | Mesenchymal | Endothelial | Epithelial | Mesenchymal |
| HALO<br>segmentation# | 5.3 | 55.2 | 39.5 | 6.7 | 56.3 | 37 |
| IMARIS<br>segmentation^ | 5.6 | 63.6 | 30.8 | 6.4 | 64 | 29.6 |

## Supplementary Information

**Figure S1.**
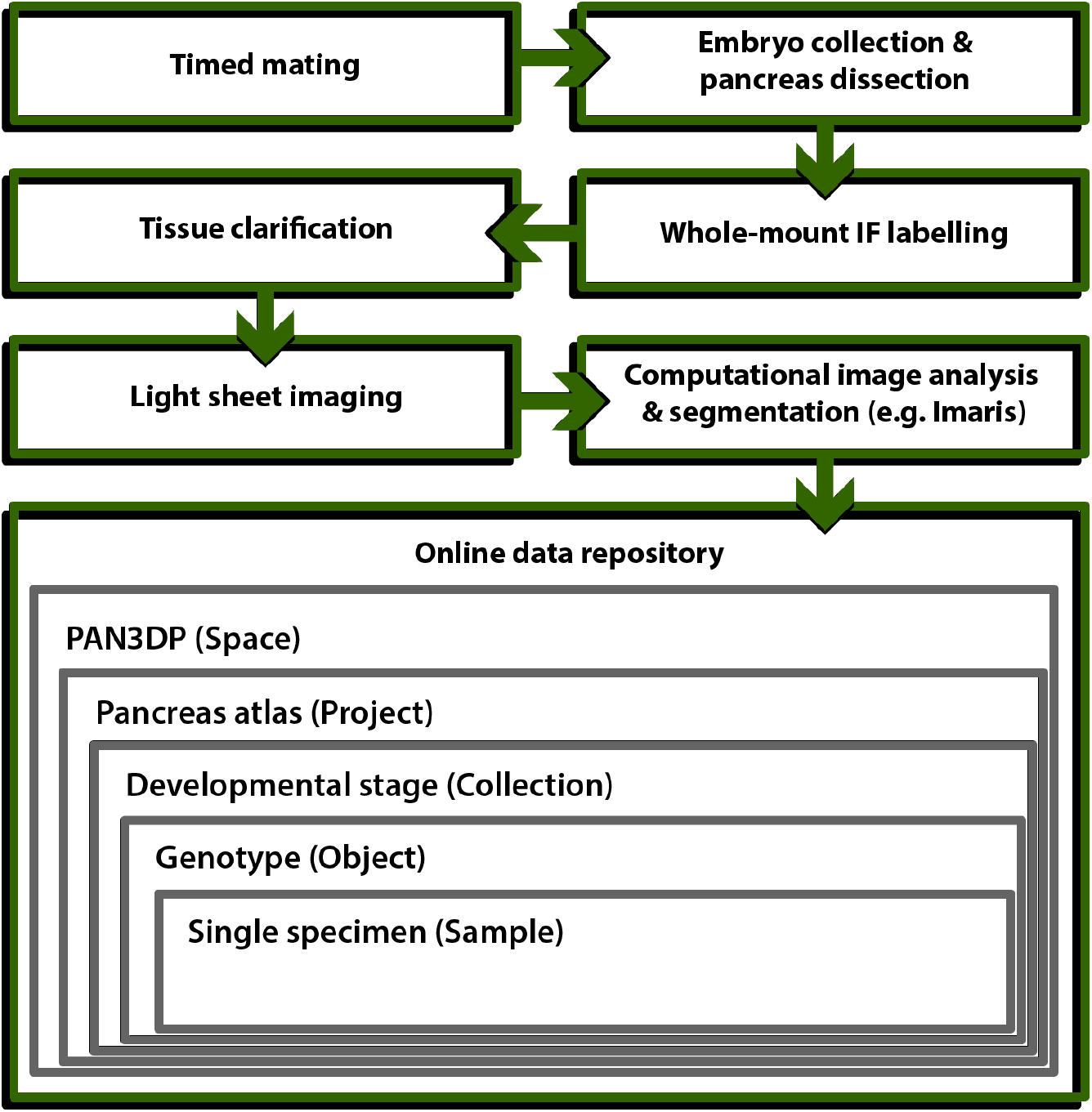
Schematic representation of data collection and deposition. Flowchart describing the experimental steps used to generate the image dataset presented here and the organizational structure of the online data repository in which they can be accessed.

**Figure S2.**
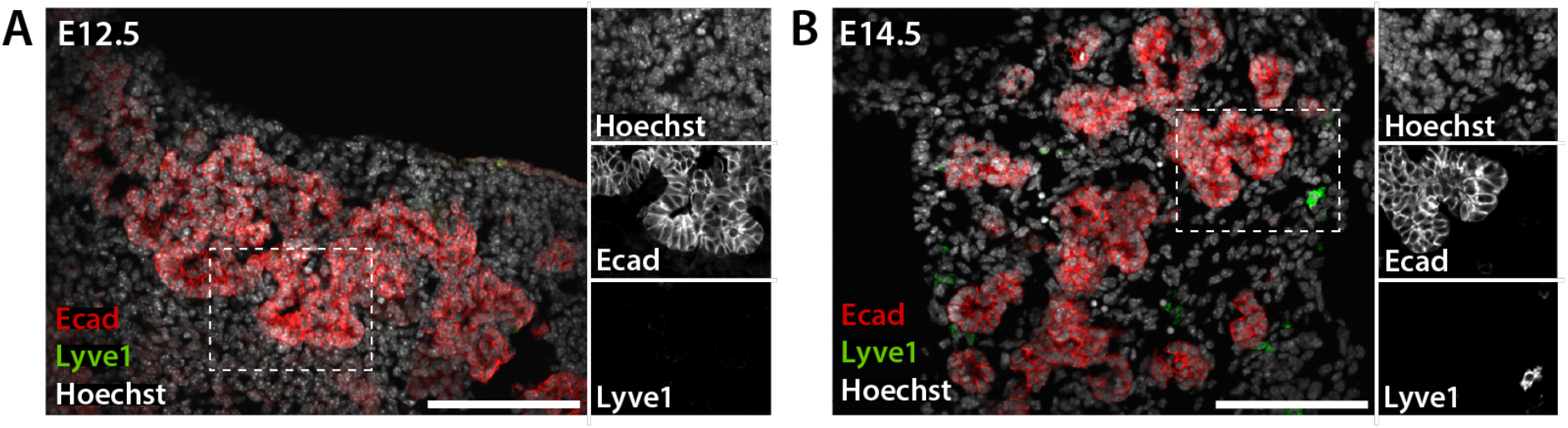
Lymphatic cells represent a rare population in the embryonic pancreas. Representative IF images of pancreatic tissue sections at E12.5 (A) and E14.5 (B). IF labelling for the Lymphatic vessel endothelial hyaluronan receptor 1 (Lyve1; green) marks rare lymphatic cells in the embryonic pancreas. Ecad (red) demarcates the pancreatic epithelium. Hoechst (grey) was used as nuclear counterstain. Insets show higher magnifications of the boxed regions as single channel configuration. Scale bars, 100 μm.

## Supplementary Tables

**Table S1.**
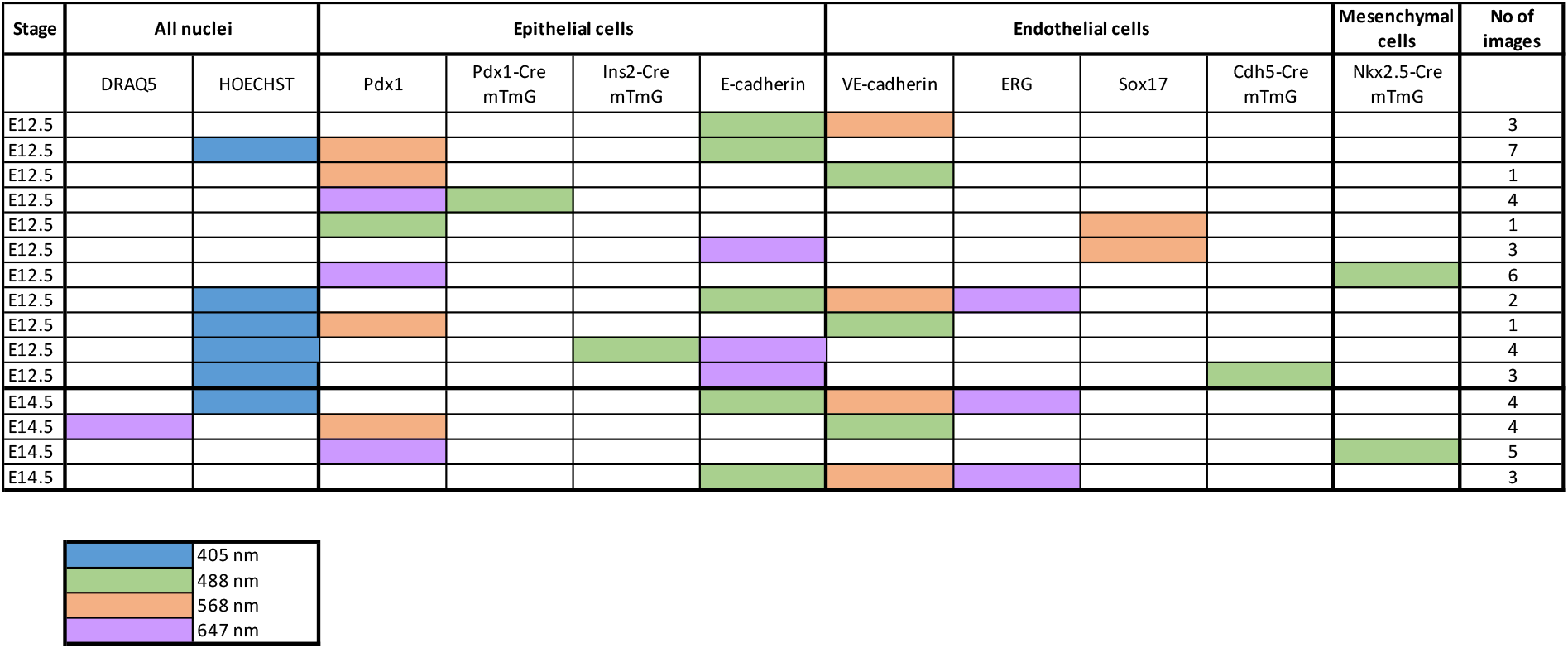
Data deposited in the Pancreas Embryonic Cell Atlas.

**Table S2.**
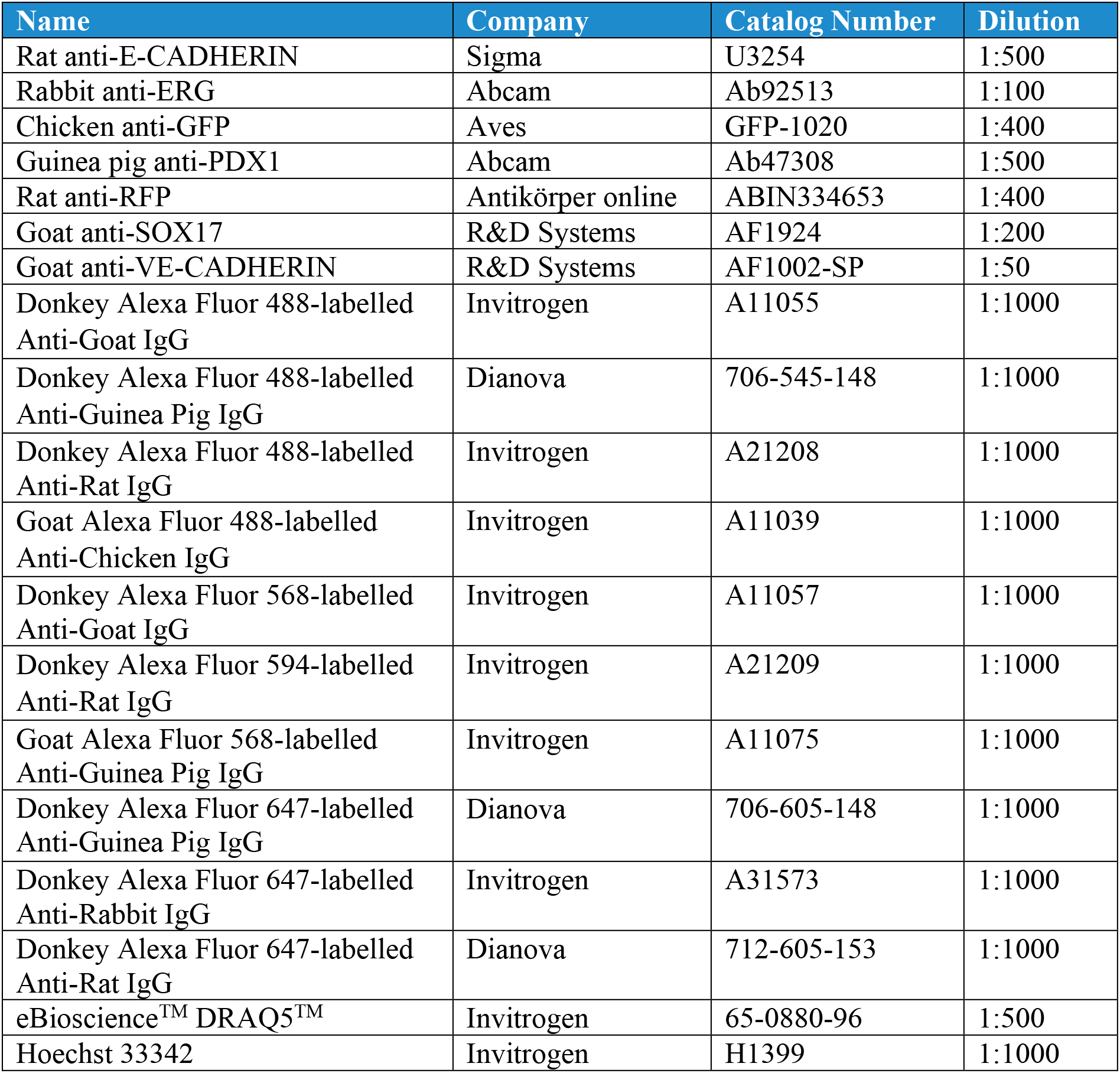
Antibodies and fluorescent dyes

## Notes

### Competing Interest Statement

The authors have declared no competing interest.

